# High genetic diversity and *bla*_NDM-1_ prevalence among *Acinetobacter baumannii* in Nigerian hospitals

**DOI:** 10.1101/2023.01.20.524999

**Authors:** Erkison Ewomazino Odih, Anderson O. Oaikhena, Anthony Underwood, Yaovi Mahuton Gildas Hounmanou, Oyinlola O. Oduyebo, Abayomi Fadeyi, Aaron O. Aboderin, Veronica O Ogunleye, Silvia Argimón, Vitus Nnaemeka Akpunonu, Phillip O. Oshun, Abiodun Egwuenu, Tochi J. Okwor, Chikwe Ihekweazu, David M. Aanensen, Anders Dalsgaard, Iruka N. Okeke

**Affiliations:** Global Health Research Unit for the Genomic Surveillance of Antimicrobial Resistance, Department of Pharmaceutical Microbiology, Faculty of Pharmacy, University of Ibadan, Ibadan, Oyo State, Nigeria; Department of Veterinary and Animal Sciences, Faculty of Health and Medical Sciences, University of Copenhagen, Copenhagen, Denmark; Centre for Genomic Pathogen Surveillance, Wellcome Sanger Institute, Cambridge, United Kingdom; Big Data Institute, University of Oxford, Oxford, United Kingdom; Department of Medical Microbiology and Parasitology, Faculty of Basic Medical Sciences, College of Medicine, University of Lagos, Lagos, Nigeria; Department of Medical Microbiology and Parasitology, University of Ilorin, Ilorin, Kwara State, Nigeria; Department of Medical Microbiology and Parasitology, Obafemi Awolowo University Teaching Hospitals Complex, Ile-Ife, Nigeria; Department of Medical Microbiology and Parasitology, University College Hospital, Ibadan, Oyo State, Nigeria; Clina-Lancet Laboratories, Victoria Island, Lagos State, Nigeria; Nigeria Centre for Disease Control, Jabi, Abuja, Nigeria

## Abstract

**Background:** *Acinetobacter baumannii* cause difficult-to-treat infections mostly among immunocompromised patients. Clinically relevant *A. baumannii* lineages and their carbapenem resistance mechanisms are sparsely described in Nigeria.

**Objective:** This study aimed to characterise the diversity and genetic mechanisms of carbapenem resistance among *A. baumannii* strains isolated from hospitals in southwestern Nigeria.

**Methods:** We sequenced the genomes of all *A. baumannii* isolates submitted to Nigeria’s antimicrobial resistance surveillance reference laboratory between 2016 – 2020 on an Illumina platform and performed *in silico* genomic characterisation. Selected strains were sequenced using the Oxford Nanopore technology to characterise the genetic context of carbapenem resistance genes.

**Results:** The 86 *A. baumannii* isolates were phylogenetically diverse and belonged to 35 distinct sequence types (STs), 16 of which were novel. Thirty-eight (44.2%) isolates belonged to none of the known international clones (ICs). Over 50% of the isolates were phenotypically resistant to 10 of 12 tested antimicrobials. Majority (n=54) of the isolates were carbapenem-resistant, particularly the IC7 (100%) and IC9 (>91.7%) strains. *bla*_OXA-23_ (34.9%) and *bla*_NDM-1_ (27.9%) were the most common carbapenem resistance genes detected. All *bla*_OXA-23_ genes were carried on Tn*2006* or Tn*2006*-like transposons. Our findings suggest that the mobilisation of a 10kb Tn*125* composite transposon is the primary means of *bla*_NDM-1_ dissemination.

**Conclusion:** Our findings highlight an increase in *bla*_NDM-1_ prevalence and the widespread transposon-facilitated dissemination of carbapenemase genes in diverse *A. baumannii* lineages in southwestern Nigeria. We make the case for improving surveillance of these pathogens in Nigeria and other understudied settings.

**Importance:** Acinetobacter baumannii are increasingly clinically relevant due to their propensity to harbour genes conferring resistance to multiple antimicrobials, as well as their ability to persist and disseminate in hospital environments and cause difficult-to-treat nosocomial infections. Little is known about the molecular epidemiology and antimicrobial resistance profiles of these organisms in Nigeria, largely due to limited capacity for their isolation, identification, and antimicrobial susceptibility testing. Our study characterised the diversity and antimicrobial resistance profiles of clinical A. baumannii in southwestern Nigeria using whole-genome sequencing. We also identified the key genetic elements facilitating the dissemination of carbapenem resistance genes within this species. This study provides key insights into the clinical burden and population dynamics of A. baumannii in hospitals in Nigeria and highlights the importance of routine whole-genome sequencing-based surveillance of this and other previously understudied pathogens in Nigeria and other similar settings.

## Background

Antimicrobial resistance (AMR) is a global public health problem, with an estimated 1.27 million deaths globally attributable to drug-resistant bacteria in 2019^1^. This estimated AMR burden is substantially higher in low-resource settings, and carbapenem-resistant *Acinetobacter baumannii* represents one of the leading causes of deaths associated with or attributable to AMR^1^.

Carbapenem resistance in *A. baumannii* is mediated primarily by carbapenemase enzymes, particularly those belonging to beta-lactamase class D (oxacillinases [OXA])^2,3^. Oxacillinases, including OXA-23, OXA-24 and OXA-58, are common among *A. baumannii* and are each encoded by various alleles with varying hydrolytic capacities. Although the intrinsic *oxaAb* gene encoding OXA-51 is harboured by all *A. baumannii* strains, only certain alleles of this gene may confer carbapenem resistance when overexpressed due to the presence of an upstream *ISAba1* promoter^4,5^. OXA-23 is the most common acquired oxacillinase in *A. baumannii*, and it is typically flanked by insertion elements that facilitate its effective spread through conjugative plasmids across different *A. baumannii* lineages^6^. The less common but significantly more potent (wider hydrolytic spectrum) New-Delhi metallo-beta-lactamases (NDM), particularly NDM-1, has been reported more frequently in recent studies, including in Nigeria and other parts of Africa^7–10^. Characterisation of the genetic diversity and contexts of carbapenemase genes in various lineages would provide insights into the population dynamics of *A. baumannii*.

Two highly successful and widely disseminated *A. baumannii* lineages, international clone (IC) 1 and IC2, predominate globally^11^. In a review of all *A. baumannii* genomes present in the National Center for Biotechnology Information’s GenBank database in 2019, 61% were IC2, and 5% were IC1^3^. International clone 2 and, to a lesser extent, IC1 strains are also responsible for majority of the carbapenem-resistant *A. baumannii* outbreaks reported in hospitals throughout the world^7,12–14^. However, as the success and global dissemination of these clones are associated with their propensity to carry multiple antimicrobial resistance determinants^13,15^, reports of increasing carbapenem resistance among *A. baumannii* isolates belonging to non-major global clones and the emergence and spread of previously uncharacterised highly resistant clones in different hospital settings is worrisome^16–18^. This necessitates the characterisation of the *A. baumannii* diversity and prevalent resistance mechanisms in different geographical contexts to inform future infection prevention and control measures and vaccination efforts. This study aimed to determine and characterise the lineages and carbapenem resistance mechanisms of *A. baumannii* isolates in southwestern Nigeria.

## Methods

### Ethical considerations

This study was approved by the University of Ibadan/University College Hospital (UI/UCH) Ethics Committee (UI/EC/19/0632). Patients were not actively recruited for this study and all associated patient data were anonymised before being retrieved for analysis.

### Isolate collection

All *Acinetobacter* isolates included in this study were isolated between 2016 – 2020 and submitted to Nigeria’s AMR surveillance reference laboratory at the University College Hospital, Ibadan, Nigeria. These isolates were collected as part of routine surveillance of WHO global priority pathogens in Nigeria and were isolated from blood, cerebrospinal fluid, and rectal swab samples. Submitting laboratories/hospitals included Lagos University Teaching Hospital, Idi-Araba, Lagos State; Clina-Lancet Laboratories, Victoria Island, Lagos State; EL-LAB Medical Diagnostics, Festac, Lagos State; Obafemi Awolowo University Teaching Hospitals Complex, Ile-Ife, Osun State; University College Hospital, Ibadan, Oyo State; University of Ilorin Teaching Hospital, Ilorin, Kwara State, and Babcock University Teaching Hospital, Ilishan-Remo, Ogun State. Where available, associated metadata on sample type, collection date, and patient hospitalisation status was obtained from the reference laboratory metadata database. All cryopreserved presumptive *Acinetobacter* isolates were resuscitated on CHROMagar™ Acinetobacter media (CHROMagar, Paris, France) and preliminarily identified using the VITEK 2 automated system (bioMérieux, Inc., Marcy-l’Étoile, France) with Gram-negative identification cards (reference number: 21341) according to the manufacturer’s instructions.

### Antimicrobial susceptibility testing

We determined the phenotypic susceptibility of the isolates to clinically relevant antimicrobials using the VITEK AST N281 cards (reference number: 414531) on the VITEK 2 automated system. The following antimicrobials were tested: cefepime, ceftazidime, ciprofloxacin, doripenem, gentamicin, imipenem, levofloxacin, meropenem, minocycline, piperacillin/tazobactam, ticarcillin/clavulanic acid, and tigecycline. The minimum inhibitory concentration (MIC) values of all tested antimicrobials except tigecycline were interpreted according to the Clinical Laboratory Standards Institute (CLSI) clinical breakpoints^19^. Tigecycline MIC values were interpreted according to EUCAST clinical breakpoints^20^ as the CLSI guidelines did not provide clinical cut-offs for tigecycline in *A. baumannii*. All interpretations were done using the AMR R package (version 1.8.1; https://msberends.github.io/AMR/).

### Genomic DNA extraction and whole-genome sequencing

We extracted genomic DNA from all the presumptively identified *Acinetobacter baumannii* complex isolates, prepared double-stranded genomic DNA libraries, and sequenced the libraries on an Illumina platform as previously described^7^. After preliminary analyses of the short-read whole-genome sequencing (WGS) data, we selected representatives of the different *A. baumannii* lineages identified in our dataset and carried out long-read whole-genome sequencing of these isolates using the Oxford Nanopore technology to obtain completely assembled genomes for comprehensive analyses. Genomic DNA was re-extracted from the selected isolates using the A&A Genomic Mini AX Bacteria+ kit (A&A Biotechnology, Gdańsk, Poland) to obtain less fragmented DNA. Long-read sequencing libraries were then generated using the Rapid Barcoding Sequencing Kit (SQK-RBK004) and sequenced on a MinION Flow Cell (R9.4.1) with MinKNOW version 22.08.9 (Oxford Nanopore Technologies, Inc., Oxford, United Kingdom). We then carried out super-accuracy base calling and demultiplexing on the generated reads using Guppy version 6.3.8.

### Whole-genome sequence analysis

We performed de-novo genome assembly, species identification, and quality control of all short-read genomes using the Global Health Research Unit (GHRU) protocol (https://www.protocols.io/view/ghru-genomic-surveillance-of-antimicrobial-resista-bpn6mmhe). All assemblies with > 300 contigs, genome sizes < 3.3 mb or > 4.7 mb, an N50 score < 25000, and containing >5% of contaminating single nucleotide variants of core genes were excluded from downstream analyses. Long-read sequences were assembled using the Trycycler pipeline^21^, and the generated circularised assemblies were then polished using Medaka v1.7.2. To generate high-quality complete assemblies, the generated long-read assemblies were then polished with the short reads using Polypolish^22^.

We performed a single nucleotide polymorphism (SNP)-based phylogenetic reconstruction analysis to determine the phylogenetic relationships between the identified *A. baumannii* strains. The raw reads of all samples were mapped to a reference sequence (Genbank accession: GCA_000830055.1) using the BWA-MEM algorithm with BWA^23^ v0.7.17, and possible duplicates were marked and removed using Picard v2.21.6 (http://broadinstitute.github.io/picard). Variant sites were called based on the alignment to the reference sequence using BCFtools^24^ v1.9, and low-quality variants were removed. Variant sites were extracted using SNP-sites^25^ v2.4.1 and concatenated into pseudogenomes for each of the samples, as well as the reference sequence, after which all pseudogenomes were combined to form a pseudoalignment. We then used RAxML-NG^26^ v1.1.0 to construct a maximum likelihood (ML) phylogeny with 50 starting trees and 1000 bootstrap replicates using the general time reversible gamma (GTR+G) model with the Lewis method for ascertainment bias correction^27^.

Multi-locus sequence types were determined from the assembled genomes using the R package MLSTar^28^ v0.1.5 with the Oxford MLST scheme^29^. The detected sequence types were assigned to IC groups if they had no more than a double locus variation from the STs in the nine defined ICs^30,31^. Lipooligosaccharide outer core loci and capsular polysaccharide loci were identified using Kaptive^32^ v2.0.3. Identified loci with at least a ‘good’ confidence match were reported. All genomes and assembly fragments were annotated using Bakta^33^ v1.5.1 with database version 4.0.0. Antimicrobial resistance genes carried by each isolate were identified using AMRfinderplus^34^ v3.10.24 with database version 2022-04-04.1. Using only the complete assemblies, the genetic contexts of carbapenem resistance genes were observed in Artemis, and genomic resistance islands were identified using the IslandViewer 4 tool^35^. Insertion sequence (IS) elements were identified using the BLAST tool on the ISfinder database (https://www-is.biotoul.fr/blast.php). Using GView Server (https://server.gview.ca/), we mapped the draft assemblies of the remaining isolates in each clone to the complete assembly of the long-read sequenced representative strain to determine representativeness.

Genomic context results for the representative strain were extrapolated for isolates that had a perfect mapping (> 95% coverage) to the representative assembly. For carbapenem-resistant strains without complete assemblies for representative sequences, we identified the contigs carrying the resistance genes and associated elements using BLAST (https://blast.ncbi.nlm.nih.gov/Blast.cgi). Genetic structure comparisons were carried out using Clinker^36^.

### Statistical analysis

All statistical analyses were carried out in R v4.2.1. Proportions were compared between groups using the Fisher’s exact test with false discovery rate correction for multiple testing. The Dunn’s test with Bonferroni correction for multiple testing was used following a Kruskal-Wallis test to carry out pairwise comparisons of the numbers of resistance determinants between multiple groups. P values less than 0.05 were considered statistically significant.

### Data availability

The raw reads of all 86 *A. baumannii* genomes have been deposited in the European Nucleotide Archive (https://www.ebi.ac.uk/ena) with study accession PRJEB29739. Accession numbers for each sample are listed in Table S1.

## Results

### Isolate collection

One hundred and twenty-five isolates were submitted as presumptive *A. baumannii* to the reference laboratory during the study period. Of these, 100 isolates were confirmed as *Acinetobacter* species based on WGS identification and included in the analyses. A further seven isolates originally submitted to the reference laboratory as *Klebsiella pneumoniae* (n=3), *Escherichia coli* (n=2), *Enterobacter cloaceae* (n=1), and *Staphylococcus haemolyticus* (n=1) but determined to be misidentified and confirmed to be *A. baumannii* by WGS were also included. Thus, a total of 107 isolates, including 36 isolates characterised in our previous study^7^, were included in the analyses. These comprised *A. baumannii* (86; 80.4%), *A. nosocomialis* (16; 15.0%), *A. haemolyticus* (two, 1.9%), *A. pittii* (one; 0.9%), *A. indicus* (one; 0.9%), and *A. variabilis* (one; 0.9%). Specimen information was unavailable for 50 isolates. Of the remaining 57 isolates, 36 were isolated from blood (33 *A. baumannii*, two *A. nosocomialis*, and one *A. variabilis*), 20 isolates were from rectal swabs (19 *A. baumannii* and one *A. pittii*), and there was one *A. baumannii* isolate from cerebrospinal fluid.

### Diversity of clinical *Acinetobacter baumannii* in southwestern Nigeria

The isolates belonged to 35 different sequence types (Oxford scheme), 16 of which were novel types (n=25; 29.1%) (Figure 1). The top 10 other sequence types detected include ST1089 (n=10; 11.6%), ST1114/1841 (n=9; 10.5%), ST229 (n=7; 8.1%), ST231 (n=6; 7.0%), ST2146 (n=4; 4.7%), ST369 (n=4; 4.7%), ST1936 (n=3; 3.5%), ST2151 (n=3; 3.5%), ST862 (n=3; 3.5%) and ST1418 (n=2; 2.3%). One isolate could not be typed as it was missing the *gdhB* locus. Nearly half (38; 44.2%) of the isolates were either singletons or belonged to none of the nine known ICs. The remaining isolates were IC2 (n=15; 17.4%), IC9 (n=13; 15.1%), IC1 (n=9; 10.5%), IC7 (n=7; 8.1%), or IC8 (n=3; 3.5%). All identified ICs formed distinct phylogenetic clusters with no geographic clustering (Figure 2). All the major lineages, except IC8 which comprised only three isolates, were detected in at least two different locations. IC7 was the only lineage detected in all five states sampled (five of the seven institutions).

**Figure 1:**
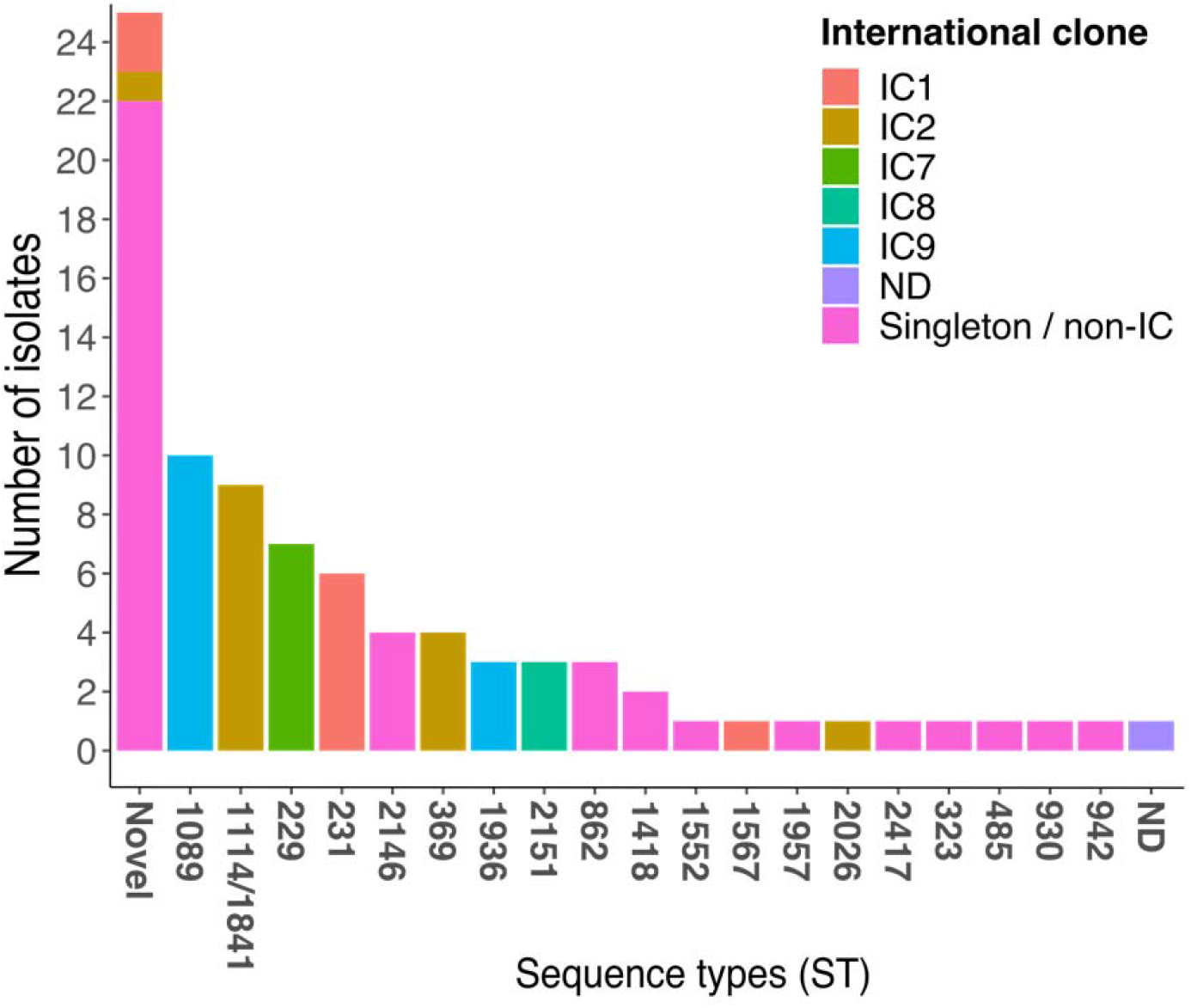
Multi-locus sequence types and international clone distribution of 86 *A. baumannii* isolates from southwestern Nigeria, 2016 – 2020.

**Figure 2:**
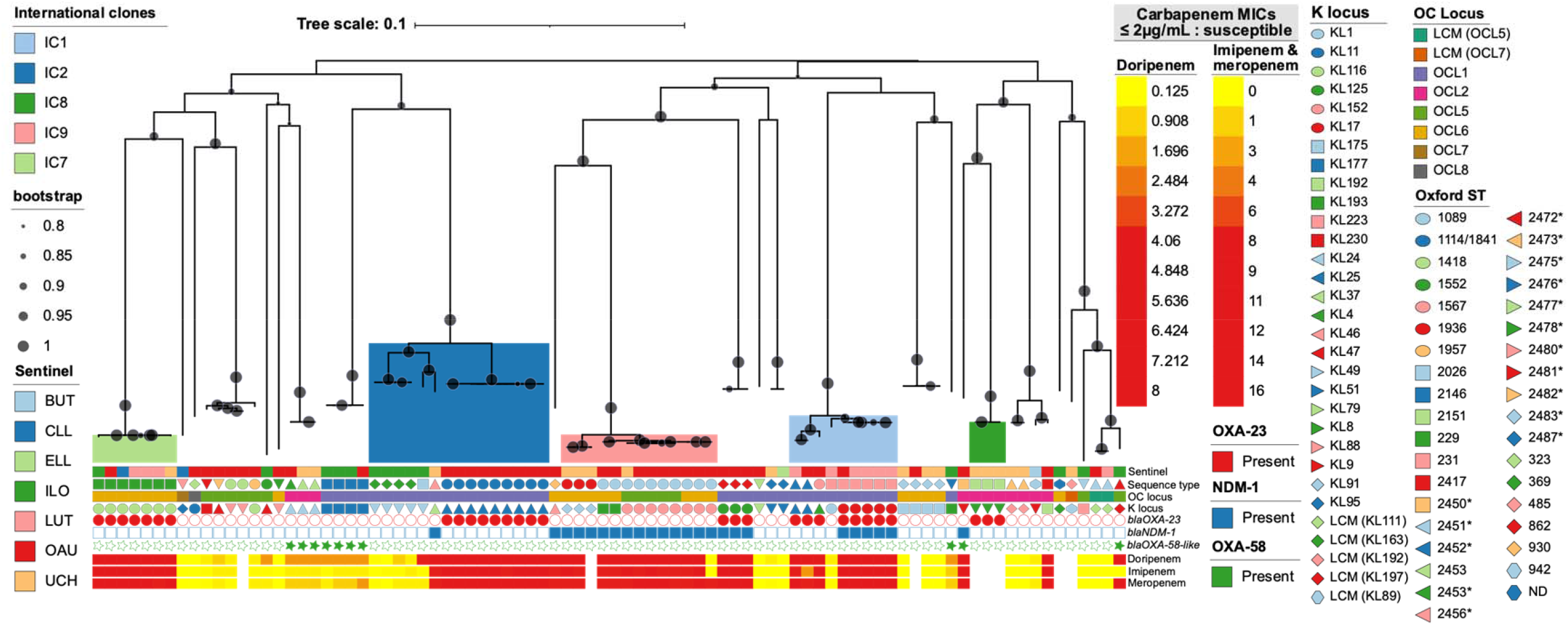
Maximum likelihood phylogeny of *A. baumannii* isolates. Coloured clades represent international clones. **BUT** – Babcock University Teaching Hospital, Ilishan-Remo, Ogun State; **CLL** – Clina-Lancet Laboratories, Victoria Island, Lagos State; **ELL** – EL-LAB Medical Diagnostics, Festac, Lagos State; **ILO** – University of Ilorin Teaching Hospital, Ilorin, Kwara State; **LUT** – Lagos University Teaching Hospital, Idi-Araba, Lagos State; **OAU** – Obafemi Awolowo University Teaching Hospitals Complex, Ile-Ife, Osun State; **UCH** – University College Hospital, Ibadan, Oyo State.

There was a high diversity with respect to the outer polysaccharide capsular loci (KL), with 31 distinct KL configurations detected among the isolates. The most common KL loci detected include KL25 (n = 11; 12.8%), KL116 (8; 9.3%), and KL152 (8; 9.3%). There was lesser diversity of the lipooligosaccharide outer core loci (OCL), but there was evidence of homologous recombination of the OCL between distantly related lineages. The majority of the isolates (35; 40.7%) had the OCL1 configuration. Twenty-two isolates (25.6%) had OCL6, 13 (15.1%) had OCL5, 11 (12.8%) had OCL2, and there was one isolate each with the OCL7 (1.2%) and OCL8 (1.2%) configuration. The remaining three isolates matched the OCL5 (n = 2; coverage 80.77%; 7/9 expected genes present) and OCL7 (n = 1; coverage 77.71%; 6/9 expected genes present) configurations most closely but with no match confidence.

### Antimicrobial resistance

Phenotypic resistance rates to all the tested antimicrobials were high among the *A. baumannii* isolates, with at least 50% resistance recorded for 10 of the 12 tested antimicrobials. Only imipenem (43%) and minocycline (23%) had lower resistance rates. The resistance rates were however significantly higher among known international clone lineages compared to the singletons/non-international clones (adjusted p ≤ 0.009; Table 1). The IC7 strains had the highest rates of resistance; all seven isolates (100%) were resistant to meropenem, imipenem, doripenem, and all the other tested antimicrobials except tigecycline (57.1%) and minocycline (0%). The IC9 strains had similarly high resistance rates to the carbapenems and other antimicrobials, as well as low resistance rates to the tetracyclines (tigecycline – 58.3%; minocycline – 16.7%). Interestingly, the globally disseminated IC1 and IC2 strains were all resistant to tigecycline (100%) but showed relatively lower resistance to the carbapenems compared to IC7 and IC9.

**Table 1:** Antimicrobial resistance rates of clinical *A. baumannii* isolates from southwestern Nigeria, 2016 – 2020.

| Antimicrobial | Lineages (% resistance) |  |  |  |  | *p* * | Adjusted *p* *** |
| --- | --- | --- | --- | --- | --- | --- | --- |
|  | IC1<br>(n = 9) | IC2<br>(n = 15) | IC7<br>(n = 7) | IC9<br>(n = 12) | Singleton/<br>non-IC<br>(n = 35) |  |  |
| Cefepime | 100 | 100 | 100 | 100 | 65.7 | 0.002 | 0.003 |
| Ceftazidime | 100 | 93.3 | 100 | 100 | 57.1 | <0.001 | 0.001 |
| Ciprofloxacin | 100 | 93.3 | 100 | 100 | 65.7 | 0.009 | 0.009 |
| Doripenem | 88.9 | 73.3 | 100 | 91.7 | 45.7 | 0.002 | 0.003 |
| Gentamicin | 100 | 100 | 100 | 100 | 57.1 | <0.001 | <0.001 |
| Imipenem | 88.9 | 66.7 | 100 | 91.7 | 20 | <0.001 | <0.001 |
| Levofloxacin | 100 | 93.3 | 100 | 100 | 65.7 | 0.009 | 0.009 |
| Meropenem | 88.9 | 66.7 | 100 | 100 | 40 | <0.001 | <0.001 |
| Minocycline | 66.7 | 73.3 | 0 | 16.7 | 11.4 | <0.001 | <0.001 |
| Piperacillin/<br>Tazobactam | 100 | 100 | 100 | 100 | 60 | <0.001 | <0.001 |
| Ticarcillin/<br>Clavulanic acid | 88.9 | 80 | 100 | 100 | 54.3 | 0.003 | 0.004 |
| Tigecycline | 100 | 100 | 57.1 | 58.3 | 51.4 | <0.001 | <0.001 |

\* Fisher's exact test comparison of resistance rates between the different lineage groups;
\*\*Adjusted p values with false discovery rate correction

Although the non-international clone isolates had significantly lower resistance rates compared to the isolates within the known ICs, they still showed at least 50% resistance to most of the tested antibiotics, including cefepime (65.7%), ciprofloxacin (65.7%), levofloxacin (65.7%), piperacillin/tazobactam (60%), ceftazidime (57.1%), gentamicin (57.1%), ticarcillin/clavulanic acid (54.3%), and tigecycline (51.4%). Resistance rates to the carbapenems were similarly lower among these strains but were nonetheless noteworthy (doripenem – 45.7%; meropenem – 40%; imipenem – 20%). Similar to the phenotypic results, the number of antibiotic classes for which specific lineages carried at least one resistance-conferring gene was significantly different. The IC7 strains carried genes conferring resistance to significantly more antimicrobial classes compared to the IC2 (mean: 10 classes vs 7 classes; adjusted p=0.01), IC8 (mean: 10 vs 3; p=0.001), and non-IC/singleton (mean: 10 vs 5; p=0.0001) strains (Figure 3). Interestingly, however, with disaggregated data, a novel ST (ST2456) carried resistance genes conferring resistance to the highest number of antimicrobial classes (median: 12 classes), followed by the IC7 strains (10 classes) and another novel ST, ST2450 (10 classes).

**Figure 3:**
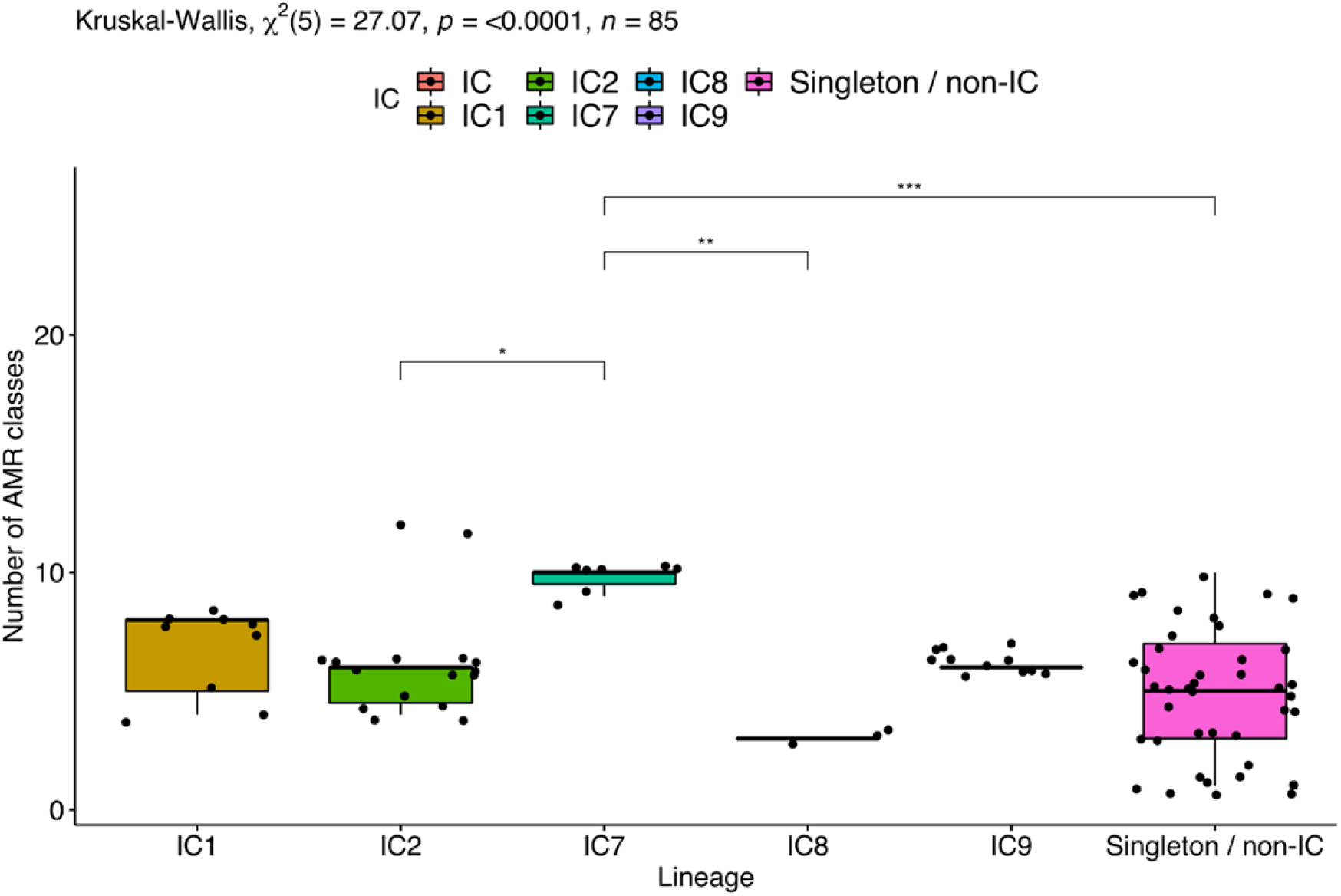
Comparison of the number of antimicrobial classes for which *A. baumannii* isolates in each lineage carried at least one resistance-conferring gene.

### Distribution and genomic context of acquired carbapenem resistance genes

The *bla*_OXA-23_ gene, present in 30 (34.9%) of the 86 isolates, was the most common acquired carbapenem resistance gene detected, followed closely by *bla*_NDM-1_ (n=24; 27.9%). The only other carbapenem resistance genes detected among the isolates were the *bla*_OXA-58_-like genes (*bla*_OXA-58_ and *bla*_OXA-420_), which were present in 10 (11.6%) isolates. The *bla*_OXA-23_ genes were almost exclusive to the international clones, being carried predominantly by IC1 (8/9), IC2 (9/15), IC7 (7/7) and IC8 (3/3) isolates; only three non-IC isolates (ST862) harboured this gene (Figure 4). These three isolates, as well as five of the IC1 isolates (ST231), co-carried the *bla*_NDM-1_ gene in addition to *bla*_OXA-23_. Among the remaining six IC2 isolates, one with a novel sequence type (ST2456) carried the *bla*_NDM-1_ gene, rather than *bla*_OXA-23_, while the other five isolates did not carry any carbapenem resistance gene. IC9 isolates (ST1089 and ST1936) were the predominant lineage carrying *bla*_NDM-1_, while *bla*_OXA-58_-like genes were only present in singleton/non-IC isolates.

**Figure 4:**
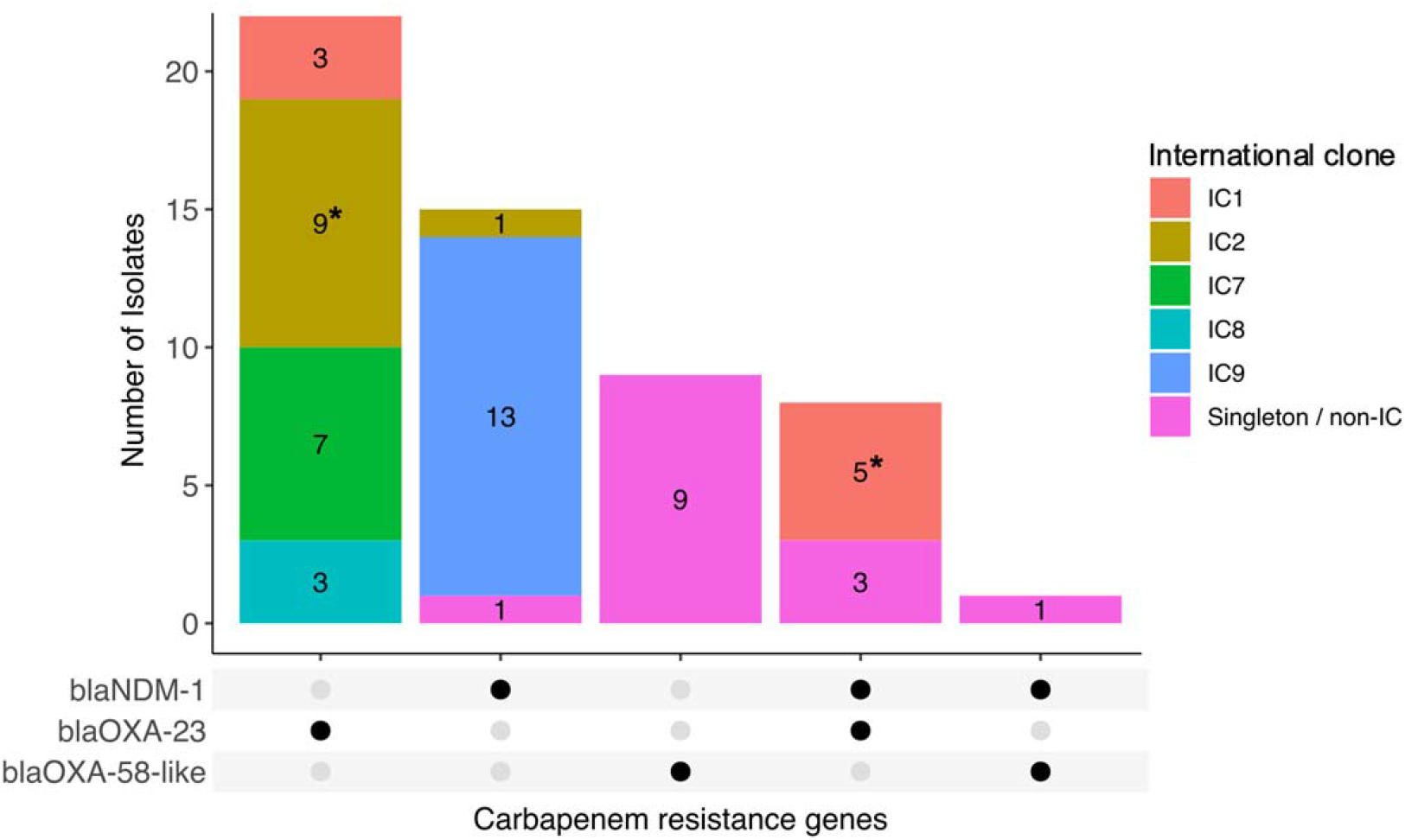
Lineage distribution and co-carriage of carbapenem resistance genes among *A. baumannii* isolates in southwestern Nigeria, 2016 – 2020. * - Carried two copies of the *bla*_OXA-23_ gene

We generated high-quality complete assemblies for representatives (five isolates) of the carbapenem-resistant lineages (Table S2) and extrapolated the genomic context results/observations to other clonal isolates. All the *bla*_OXA-23_ genes in the different clones in this study had similar immediate genetic contexts; they were all carried on a Tn*2006* transposon flanked by IS*4* family *ISAba1* inverted repeat elements, or a similar Tn*2006*-like transposon that was missing the truncated DEAD helicase gene (Figure 5). The nine ST1114/1841 isolates carried two distant (~1.5 MB apart) copies of the *bla*_OXA-23_ gene on the chromosome, each flanked by *ISAba1*. Interestingly, one of these copies was inserted just upstream of the intrinsic *bla*_OXA-51_-*like* gene (*bla*_OXA-66_). Among the five ST231 (IC1) isolates, the *bla*_OXA-23_ carbapenem resistance gene was also chromosomally located. Like the ST1114/1841 isolates, these isolates also had two copies of the *bla*_OXA-23_ gene, both proximal, flanked by *ISAba1*, and inserted into the chromosome without being associated with any other mobile elements; one copy was inserted between the *ycfP* and *menH* genes. There was only a single copy of the *bla*_OXA-23_ gene in the seven IC7 (ST229) isolates, which was also present on the chromosome and, interestingly, located within an AbaR4-type genomic island inserted within the *yifB* (*comM* sub-family) gene.

**Figure 5:**
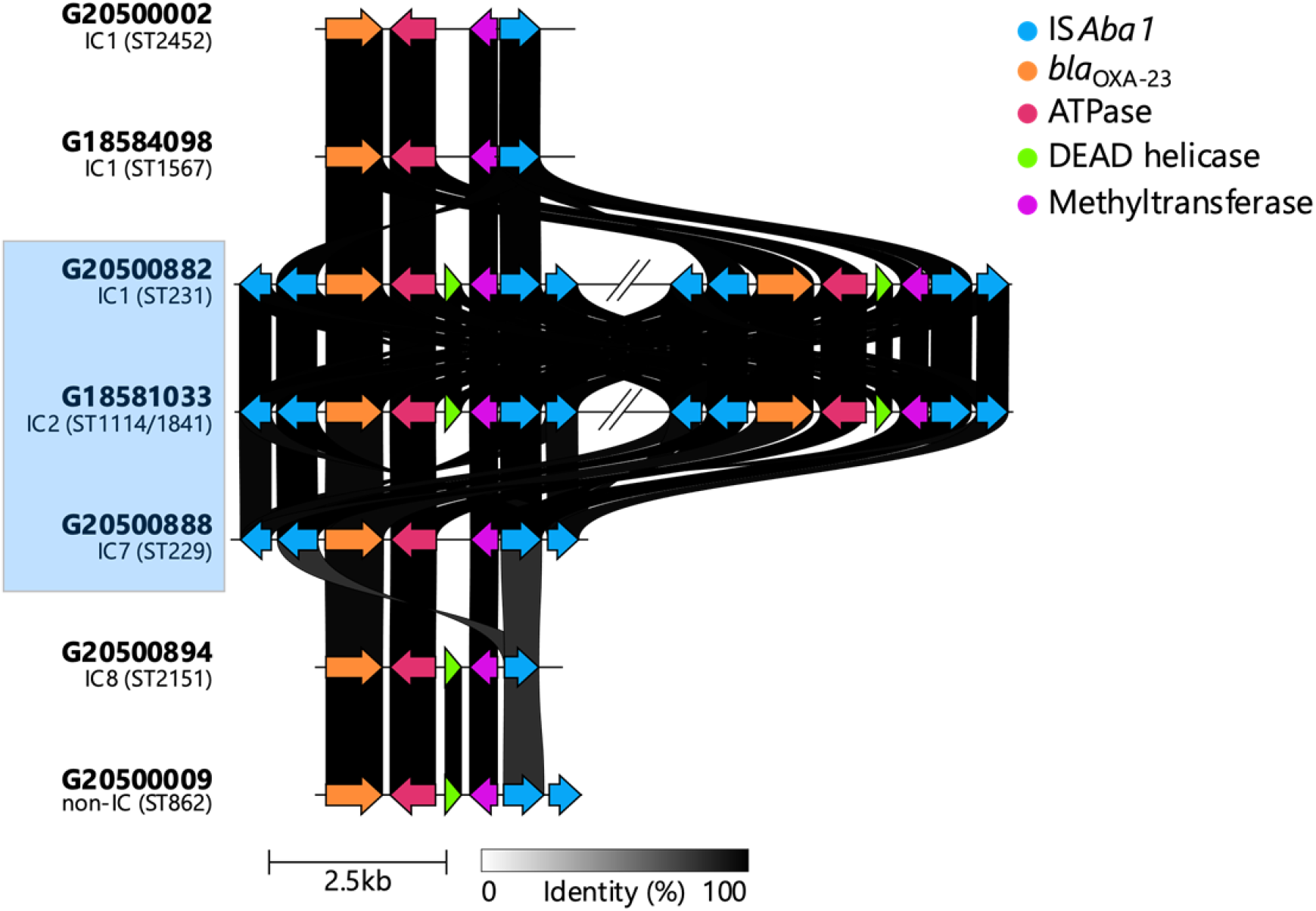
Tn*2006* and Tn*2006*-like transposons carrying the *bla*_OXA-23_ carbapenemase gene in *A. baumannii* isolates in southwestern Nigeria, 2016 – 2020. A complete IS*Aba1* unit comprises two open reading frames, both indicated with blue arrows. The two Tn*2006* copies in the ST231 (~110 kb apart) and ST1114/1841 (~1.5 mb apart) isolates are shown. Genetic contexts for lineages highlighted in the light blue box were identified based on annotated complete assemblies while the contexts for unshaded lineages were identified based on identification and annotation of the contig carrying the *bla*_OXA-23_ gene, hence the incomplete repeat element flanks.

As with both copies of the *bla*_OXA-23_ genes carried by the five ST231 (IC1) isolates, the *bla*_NDM-1_ gene was also chromosomally located and was carried on a 10kb Tn*125* composite transposon flanked by the *ISAba125* element (Figure 6). This composite transposon was inserted within a larger 25kb transposon flanked by an IS*6* family insertion sequence, IS*1006*, which also carried the *aph(6)-Id* and *aph(3”)Ib* genes. The 13 isolates belonging to IC9 (ST1089 and ST1936) had a different NDM-1 context. In these isolates, the *bla*_NDM-1_ gene was also located on the chromosome but was associated with an upstream IS*Aba125* element and located within a 7.9kb Tn*7382* transposon that also carried *aph(3’)-VI* and had two flanking IS*3* family *ISAba14* direct repeats. The remaining six *bla*_NDM-1_-carrying isolates carried the gene on a Tn*125* composite transposon like the IC1 isolates, but we could not determine if this transposon was on the chromosome or a plasmid as there was no representative complete assembly sequenced for this set.

**Figure 6:**
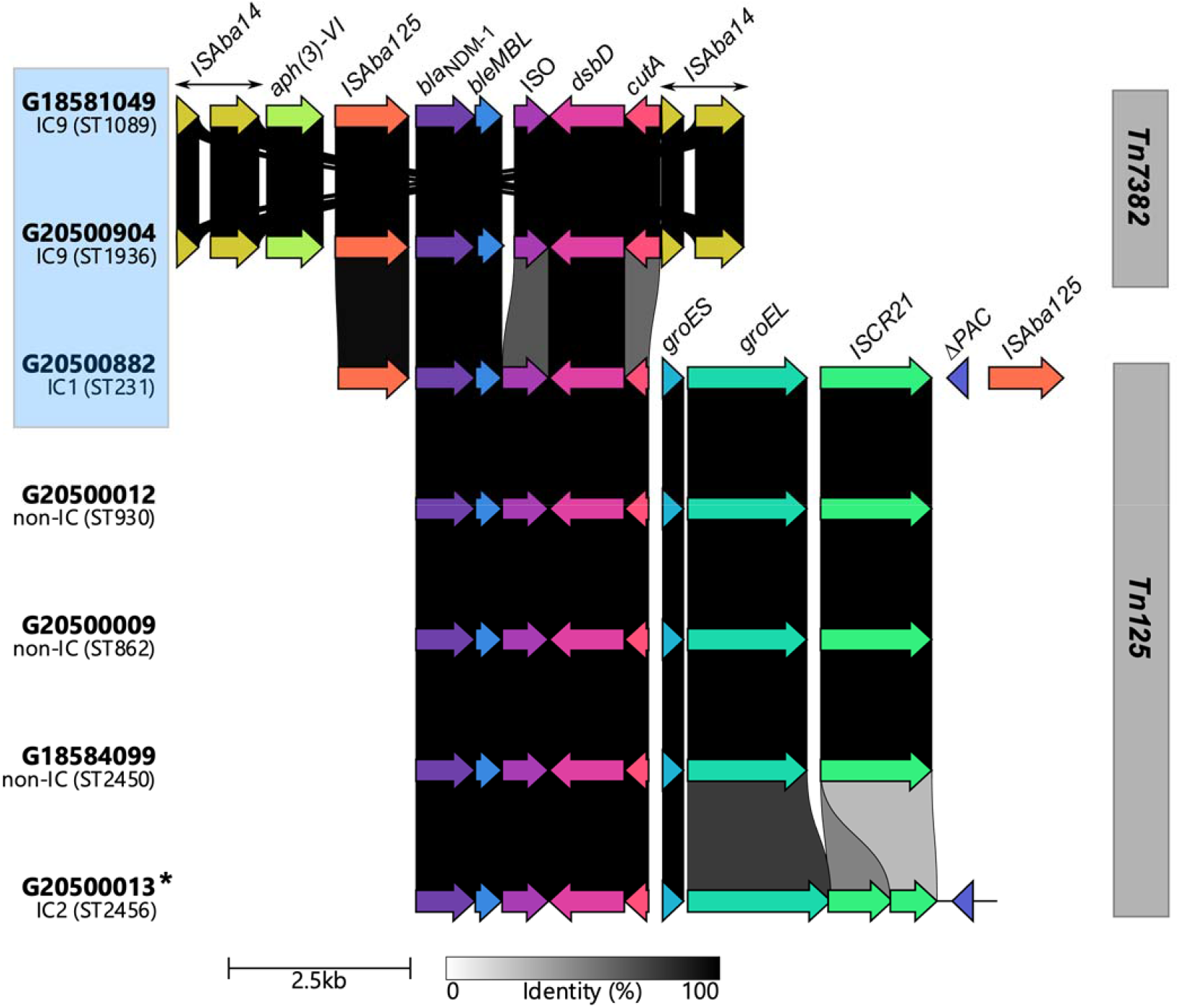
Tn*125* and Tn*7382* transposons carrying the *bla*_NDM-1_ carbapenemase gene in *A. baumannii* isolates in southwestern Nigeria, 2016 – 2020. Genetic contexts for lineages highlighted in the light blue box were identified based on annotated complete assemblies while the contexts for the other lineages were identified based on identification and annotation of the contig carrying the *bla*_NDM-1_ gene, hence the incomplete repeat element flanks. * – Split across two contigs

## Discussion

*Acinetobacter baumannii* infections remain a global public health problem, and much remains to be understood about the population structure of this species, especially in understudied settings. In this study, we discovered a large diversity of *A. baumannii* in the clinical setting of southwestern Nigeria hospitals, with 35 different sequence types, 16 of which were novel. Studies conducted in previously uncharacterised settings have reported similar findings^7,37^. A 2018 retrospective study in Colombia found seven novel STs out of the 11 detected that had been circulating for over 8 years^16^. Given the relatively small number of samples in our study, the observed high diversity and lack of phylogeographic clustering may also be indicative of widespread dissemination and underreporting of *A. baumannii* infections in Nigeria. Clinical microbiology diagnostics remains a challenge in Nigeria and most developing countries. Very few patients are cultured at all and *A. baumannii* are particularly difficult to identify using conventional diagnostics^7,38,39^. The institution of genomic surveillance for AMR in Nigeria has been coupled with efforts to improve basic microbiology capacity, and it is expected that these measures will help plug the existing gaps in the diagnostics and surveillance of these pathogens of major public health importance^40^.

This study adds to our understanding that certain clones (ICs 1-9) predominate globally and account for a large proportion of the antimicrobial resistance problem in *A. baumannii*. Nonetheless, we furthermore show that novel and emerging clones are also important in endemic settings, as evidenced by the relatively high resistance rates (≥50% resistance to eight of the 12 antimicrobials tested) of these novel STs/non-international clones, as well as their possession of carbapenem resistance determinants with identical genetic contexts to those present in the international clones. Given that carbapenem resistance contributes to clonal expansion and successful dissemination among *A. baumannii*, the local emergence of a wide variety of carbapenem-resistant variants is a noteworthy trend^15,41^. Moreover, within the known international clones, IC1 and IC2 are still regarded as the most important *A. baumannii* lineages causing infections and hospital outbreaks globally^3,42–44^. Our study however shows an increasing local prevalence of isolates belonging to the recently classified IC9 lineage relative to the globally prevalent IC2 lineage. Even more notable is the associated increased prevalence of the previously rare *bla*_NDM-1_ carbapenem resistance gene. STs belonging to IC9, mostly ST1089 (Pasteur ST85), and most of which also carry the *bla*_NDM-1_ gene or other variants such as *NDM-6*, have been reported more frequently predominantly in Africa and the Middle East^45–51^, but also in many other parts of the world^17,52–54^.

Previous studies have highlighted *bla*_OXA-23_ as the predominant carbapenem resistance gene among *A. baumannii* and shown that *bla*_NDM-1_ is rare among *A. baumannii*^55^. Our findings, however, show that although OXA-23 carbapenemases are the most common enzymes among *A. baumannii* in southwestern Nigeria, NDM-1 prevalence is also notably high in our setting and appears to be spreading between different lineages. The Tn*125* composite transposon carrying the *bla*_NDM-1_ gene in the five ST231 (IC1) isolates and six other isolates with distinct STs (ST862, ST930, ST2450, and ST2456 (IC2)) is identical in structure and composition to that previously described in an *A. baumannii* strain isolated in Germany in 2007 and other subsequent studies^55,56^. This *bla*_NDM-1_-carrying Tn*125* transposon, which is believed to have originated in *A. baumannii*, has also been demonstrated to be very frequently and efficiently mobilised, facilitating its dissemination within and between *A. baumannii* and other Enterobacterales^57^. BLAST searches of this transposon against the National Center for Biotechnology Information’s non-redundant nucleotide database revealed the presence of this transposon and its derivatives predominantly among *A. baumannii* and non-*baumannii Acinetobacter* species, but also in different plasmids and chromosomes of *E. coli*, *K. pneumoniae*, and other Enterobacterales. The *bla*_NDM-1_ context among the IC9 however had a different context – a truncated Tn*125* transposon captured within flanking *ISAba14* direct repeats and recently named Tn*7382*^58^. This transposon is only sparsely described in literature but is predominantly found among *A. baumannii* strains belonging to IC9^54,58^. The increased NDM-1 prevalence in our study and potential for intra- and inter-species spread has notable implications as this enzyme is very potent and has a wider spectrum of hydrolysis for beta-lactams and carbapenems compared to OXA-23 and OXA-58, thus grossly limiting the already limited number of treatment options for *A. baumannii* infections^59–62^.

Although all the *bla*_OXA-23_ genes in our study had similar contexts, these Tn*2006* and Tn*2006*-like transposon structures were found in diverse chromosomal backgrounds, thus adding to previous knowledge of their rapid and frequent genome mobility^6^. One interesting observation was the presence of two copies of the *bla*_OXA-23_ gene in the ST1114/1841 and ST231 strains. A previous study by Zhang and colleagues reported the duplication of a plasmid-borne *bla*_OXA-23_ gene in an *A. baumannii* strain in the presence of sub-inhibitory concentrations of carbapenem^63^. This duplication was reported to confer improved fitness in carbapenem-containing media but was also associated with a fitness cost in antibiotic-free media. Our findings, however, indicate the maintenance of multiple *bla*_OXA-23_ copies in the chromosomes of multiple strains in distinct lineages, despite the reported fitness costs and resulting instability. This is an important finding as it suggests a key evolutionary adaptation that could lead to the increased potency of OXA-23 carbapenemases, expansion of *A. baumannii* populations harbouring *bla*_OXA-23_, as well as an increased risk of mobilisation and onward dissemination of the gene.

## Conclusion

*Acinetobacter baumannii* in the hospital setting in southwestern Nigeria are highly phylogenetically diverse, highly resistant to antimicrobials, and may be underreported, indicating the urgent need to improve diagnostic capacity for and surveillance of *A. baumannii* infections both in Nigeria and other understudied settings. Our findings also suggest that there is frequent dissemination of carbapenem resistance genes between the different *A. baumannii* lineages, as well as integration and possible maintenance of these genes in the chromosomes. More local studies are needed to characterise the hospital burden of *A. baumannii* infection in Nigeria and identify contributors to environmental and clinical spread.

## Acknowledgements

We thank Ayorinde Afolayan, Gabriel Temitope Sunmonu, and Gitte Petersen for technical assistance. We also acknowledge Nonyelum Osuagwu for coordinating the collection of isolates from facilities (Clina-Lancet Laboratories and EL-LAB Medical Diagnostics) that are not originally part of Nigeria’s AMR surveillance system.

## Supplementary material legends

Table S1: Accession numbers and epidemiological information of clinical A. baumannii isolates in southwestern Nigeria, 2016 – 2020.

Table S2: Quality metrics of genome assemblies generated using long reads

## References

1. Murray, C. J. et al. Global burden of bacterial antimicrobial resistance in 2019: a systematic analysis. The Lancet 399, 629–655 (2022).

2. Peleg, A. Y., Seifert, H. & Paterson, D. L. Acinetobacter baumannii: Emergence of a Successful Pathogen. Clin. Microbiol. Rev. 21, 538–582 (2008).

3. Hamidian, M. & Nigro, S. J. Emergence, molecular mechanisms and global spread of carbapenem-resistant Acinetobacter baumannii. Microb. Genomics 5, (2019).

4. Nigro, S. J. & Hall, R. M. Does the intrinsic oxaAb (blaOXA-51-like) gene of Acinetobacter baumannii confer resistance to carbapenems when activated by ISAba1? J. Antimicrob. Chemother. 73, 3518–3520 (2018).

5. Turton, J. F. et al. The role of ISAba1 in expression of OXA carbapenemase genes in Acinetobacter baumannii. FEMS Microbiol. Lett. 258, 72–77 (2006).

6. Nigro, S. J. & Hall, R. M. Structure and context of Acinetobacter transposons carrying the oxa23 carbapenemase gene. J. Antimicrob. Chemother. 71, 1135–1147 (2016).

7. Odih, E. E. et al. Rectal Colonization and Nosocomial Transmission of Carbapenem-Resistant Acinetobacter baumannii in an Intensive Care Unit, Southwest Nigeria. Front. Med. 9, (2022).

8. Pogue, J. M., Mann, T., Barber, K. E. & Kaye, K. S. Carbapenem-resistant Acinetobacter baumannii: epidemiology, surveillance and management. Expert Rev. Anti Infect. Ther. 11, 383–393 (2013).

9. Wang, J. et al. Multidrug-resistant Acinetobacter baumannii strains with NDM-1: Molecular characterization and in vitro efficacy of meropenem-based combinations. Exp. Ther. Med. 18, 2924–2932 (2019).

10. Yehouenou, C. et al. First detection of a plasmid-encoded New-Delhi metallo-beta-lactamase-1 (NDM-1) producing Acinetobacter baumannii using whole genome sequencing, isolated in a clinical setting in Benin. Ann. Clin. Microbiol. Antimicrob. 20, 5 (2021).

11. Holt, K. et al. Five decades of genome evolution in the globally distributed, extensively antibiotic-resistant Acinetobacter baumannii global clone 1. Microb. Genomics 2, e000052 (2016).

12. Warner, W. A. et al. Molecular characterization and antimicrobial susceptibility of Acinetobacter baumannii isolates obtained from two hospital outbreaks in Los Angeles County, California, USA. BMC Infect. Dis. 16, 194 (2016).

13. Matsui, M. et al. Distribution and Molecular Characterization of Acinetobacter baumannii International Clone II Lineage in Japan. Antimicrob. Agents Chemother. 62, e02190–17 (2018).

14. Camargo, C. H. et al. Clonal spread of ArmA-and OXA-23-coproducing Acinetobacter baumannii International Clone 2 in Brazil during the first wave of the COVID-19 pandemic. J. Med. Microbiol. 71, (2022).

15. Zarrilli, R., Pournaras, S., Giannouli, M. & Tsakris, A. Global evolution of multidrug-resistant Acinetobacter baumannii clonal lineages. Int. J. Antimicrob. Agents 41, 11–19 (2013).

16. Correa, A. et al. Distinct Genetic Diversity of Carbapenem-Resistant Acinetobacter baumannii from Colombian Hospitals. Microb. Drug Resist. 24, 48–54 (2018).

17. Shrestha, S. et al. Molecular epidemiology of multidrug-resistant Acinetobacter baumannii isolates in a university hospital in Nepal reveals the emergence of a novel epidemic clonal lineage. Int. J. Antimicrob. Agents 46, 526–531 (2015).

18. Hsieh, Y.-C. et al. An Outbreak of tet(X6)-Carrying Tigecycline-Resistant Acinetobacter baumannii Isolates with a New Capsular Type at a Hospital in Taiwan. Antibiotics 10, 1239 (2021).

19. CLSI. M100Ed32 / Performance Standards for Antimicrobial Susceptibility Testing, 32nd Edition. (Wayne, PA: Clinical and Laboratory Standards Institute, 2021).

20. The European Committee on Antimicrobial Susceptibility Testing. Breakpoint tables for interpretation of MICs and zone diameters v11.0. https://www.eucast.org/clinical_breakpoints/.

21. Wick, R. R. et al. Trycycler: consensus long-read assemblies for bacterial genomes. Genome Biol. 22, 266 (2021).

22. Wick, R. R. & Holt, K. E. Polypolish: Short-read polishing of long-read bacterial genome assemblies. PLOS Comput. Biol. 18, e1009802 (2022).

23. Li, H. & Durbin, R. Fast and accurate short read alignment with Burrows-Wheeler transform. Bioinforma. Oxf. Engl. 25, 1754–1760 (2009).

24. Danecek, P. et al. Twelve years of SAMtools and BCFtools. GigaScience 10, giab008 (2021).

25. Page, A. J. et al. SNP-sites: rapid efficient extraction of SNPs from multi-FASTA alignments. Microb. Genomics 2, e000056 (2016).

26. Kozlov, A. M., Darriba, D., Flouri, T., Morel, B. & Stamatakis, A. RAxML-NG: a fast, scalable and user-friendly tool for maximum likelihood phylogenetic inference. Bioinformatics 35, 4453–4455 (2019).

27. Lewis, P. O. A Likelihood Approach to Estimating Phylogeny from Discrete Morphological Character Data. Syst. Biol. 50, 913–925 (2001).

28. Ferrés, I. & Iraola, G. MLSTar: automatic multilocus sequence typing of bacterial genomes in R. PeerJ 6, e5098 (2018).

29. Bartual, S. G. et al. Development of a Multilocus Sequence Typing Scheme for Characterization of Clinical Isolates of Acinetobacter baumannii. J. Clin. Microbiol. 43, 4382–4390 (2005).

30. Higgins, P. G., Prior, K., Harmsen, D. & Seifert, H. Development and evaluation of a core genome multilocus typing scheme for whole-genome sequence-based typing of Acinetobacter baumannii. PLOS ONE 12, e0179228 (2017).

31. Xanthopoulou, K. et al. First Report of New Delhi Metallo-β-Lactamase-6 (NDM-6) in a Clinical Acinetobacter baumannii Isolate From Northern Spain. Front. Microbiol. 11, (2020).

32. Lam, M. M. C., Wick, R. R., Judd, L. M., Holt, K. E. & Wyres, K. L. Y. 2022. Kaptive 2.0: updated capsule and lipopolysaccharide locus typing for the Klebsiella pneumoniae species complex. Microb. Genomics 8, 000800.

33. Schwengers, O. et al. Bakta: rapid and standardized annotation of bacterial genomes via alignment-free sequence identification. Microb. Genomics 7, 000685 (2021).

34. Feldgarden, M. et al. AMRFinderPlus and the Reference Gene Catalog facilitate examination of the genomic links among antimicrobial resistance, stress response, and virulence. Sci. Rep. 11, 12728 (2021).

35. Bertelli, C. et al. Islandviewer 4: expanded prediction of genomic islands for larger-scale datasets. Nucleic Acids Res. 45, W30–W35 (2017).

36. Gilchrist, C. L. M. & Chooi, Y.-H. clinker & clustermap.js: automatic generation of gene cluster comparison figures. Bioinformatics 37, 2473–2475 (2021).

37. Opazo-Capurro, A. et al. Evolutionary dynamics of carbapenem-resistant Acinetobacter baumannii circulating in Chilean hospitals. Infect. Genet. Evol. 73, 93–97 (2019).

38. Abubakar, I. et al. The Lancet Nigeria Commission: investing in health and the future of the nation. The Lancet 399, 1155–1200 (2022).

39. Afolayan, A. O. et al. Clones and Clusters of Antimicrobial-Resistant Klebsiella From Southwestern Nigeria. Clin. Infect. Dis. Off. Publ. Infect. Dis. Soc. Am. 73, S308–S315 (2021).

40. Okeke, I. N. et al. Establishing a national reference laboratory for antimicrobial resistance using a whole-genome sequencing framework: Nigeria’s experience. Microbiology 168, 001208.

41. Jun, S. H. et al. Clonal change of carbapenem-resistant Acinetobacter baumannii isolates in a Korean hospital. Infect. Genet. Evol. J. Mol. Epidemiol. Evol. Genet. Infect. Dis. 93, 104935 (2021).

42. Karah, N., Khalid, F., Wai, S. N., Uhlin, B. E. & Ahmad, I. Molecular epidemiology and antimicrobial resistance features of Acinetobacter baumannii clinical isolates from Pakistan. Ann. Clin. Microbiol. Antimicrob. 19, 2 (2020).

43. Khuntayaporn, P. et al. Predominance of international clone 2 multidrug-resistant Acinetobacter baumannii clinical isolates in Thailand: a nationwide study. Ann. Clin. Microbiol. Antimicrob. 20, 19 (2021).

44. Wohlfarth, E. et al. The evolution of carbapenem resistance determinants and major epidemiological lineages among carbapenem-resistant Acinetobacter baumannii isolates in Germany, 2010-2019. Int. J. Antimicrob. Agents 106689 (2022) doi:10.1016/j.ijantimicag.2022.106689.

45. Al-Hassan, L. et al. Molecular Epidemiology of Carbapenem-Resistant Acinetobacter baumannii From Khartoum State, Sudan. Front. Microbiol. 12, (2021).

46. Zafer, M. M., Hussein, A. F. A., Al-Agamy, M. H., Radwan, H. H. & Hamed, S. M. Genomic Characterization of Extensively Drug-Resistant NDM-Producing Acinetobacter baumannii Clinical Isolates With the Emergence of Novel bla ADC-257. Front. Microbiol. 12, 736982 (2021).

47. Uwingabiye, J. et al. Clonal diversity and detection of carbapenem resistance encoding genes among multidrug-resistant Acinetobacter baumannii isolates recovered from patients and environment in two intensive care units in a Moroccan hospital. Antimicrob. Resist. Infect. Control 6, 99 (2017).

48. Salloum, T. et al. Genomic mapping of ST85 blaNDM-1 and blaOXA-94 producing Acinetobacter baumannii isolates from Syrian Civil War Victims. Int. J. Infect. Dis. 74, 100–108 (2018).

49. Jaidane, N. et al. Whole-genome sequencing of NDM-1-producing ST85 Acinetobacter baumannii isolates from Tunisia. Int. J. Antimicrob. Agents 52, 916–921 (2018).

50. Maamar, E. et al. NDM-1-and OXA-23-producing Acinetobacter baumannii isolated from intensive care unit patients in Tunisia. Int. J. Antimicrob. Agents 52, 910–915 (2018).

51. Higgins, P. G. et al. Molecular Epidemiology of Carbapenem-Resistant Acinetobacter baumannii Isolates from Northern Africa and the Middle East. Antibiot. Basel Switz. 10, 291 (2021).

52. Higgins, P. G. et al. Molecular Epidemiology of Carbapenem-Resistant Acinetobacter baumannii Strains Isolated at the German Military Field Laboratory in Mazar-e Sharif, Afghanistan. Microorganisms 9, 2229 (2021).

53. Heydari, F., Mammina, C. & Koksal, F. 2015. NDM-1-producing Acinetobacter baumannii ST85 now in Turkey, including one isolate from a Syrian refugee. J. Med. Microbiol. 64, 1027–1029.

54. Fernández-Cuenca, F. et al. First identification of blaNDM-1 carbapenemase in blaOXA-94-producing Acinetobacter baumannii ST85 in Spain. Enfermedades Infecc. Microbiol. Clínica 38, 11–15 (2020).

55. Pfeifer, Y. et al. Molecular characterization of blaNDM-1 in an Acinetobacter baumannii strain isolated in Germany in 2007. J. Antimicrob. Chemother. 66, 1998–2001 (2011).

56. Poirel, L. et al. Tn125-Related Acquisition of blaNDM-Like Genes in Acinetobacter baumannii. Antimicrob. Agents Chemother. 56, 1087–1089 (2012).

57. Bontron, S., Nordmann, P. & Poirel, L. Transposition of Tn125 Encoding the NDM-1 Carbapenemase in Acinetobacter baumannii. Antimicrob. Agents Chemother. 60, 7245–7251 (2016).

58. Hamed, S. M., Hussein, A. F. A., Al-Agamy, M. H., Radwan, H. H. & Zafer, M. M. Tn7382, a novel composite transposon harboring blaNDM-1 and aphA6 in Acinetobacter baumannii. J. Glob. Antimicrob. Resist. 30, 414–417 (2022).

59. Fishbain, J. & Peleg, A. Y. Treatment of Acinetobacter Infections. Clin. Infect. Dis. 51, 79–84 (2010).

60. Isler, B., Doi, Y., Bonomo, R. A. & Paterson, D. L. New Treatment Options against Carbapenem-Resistant Acinetobacter baumannii Infections. Antimicrob. Agents Chemother. 63, e01110–18 (2019).

61. O’Donnell, J. N., Putra, V. & Lodise, T. P. Treatment of patients with serious infections due to carbapenem-resistant Acinetobacter baumannii: How viable are the current options? Pharmacother. J. Hum. Pharmacol. Drug Ther. 41, 762–780 (2021).

62. Zhou, S. et al. “Roar” of blaNDM-1 and “silence” of blaOXA-58 co-exist in Acinetobacter pittii. Sci. Rep. 5, 8976 (2015).

63. Zhang, L. et al. Co-evolutionary adaptations of Acinetobacter baumannii and a clinical carbapenemase-encoding plasmid during carbapenem exposure. Evol. Appl. 15, 1045–1061 (2022).

